# Geno-pheno characterization of crop rhizospheres: An integrated Raman spectroscopy and microbiome approach in conventional and organic agriculture

**DOI:** 10.1101/2025.06.10.658954

**Authors:** Yejin Son, Peisheng He, Mathew Baldwin, Guangyu Li, Zijian Wang, April Z. Gu, Jenny Kao-Kniffin

**Affiliations:** School of Integrative Plant Science, Cornell University, Ithaca, NY; School of Civil and Environmental Engineering, Cornell University, Ithaca, NY; School of Biological and Environmental Engineering, Cornell University, Ithaca, NY

**Author notes:** Corresponding Author: Jenny Kao-Kniffin.

**Keywords:** Soil microorganisms, organic agriculture, conventional agriculture, agricultural microbiome, Raman single cell spectroscopy, food security

## Abstract

1

In this study, we examined phenotypic and compositional patterns in rhizosphere microbial communities across conventionally and organically managed farms to assess impacts on soil microbiomes. We employed newly developed single-cell Raman microspectroscopy (SCRS)-based community phenotypic profiling analysis with microbiome 16S rRNA gene amplicon sequencing to compare the soil microbial communities of alfalfa, carrot, corn, lettuce, potato, soybean, squash, tomato, triticale, wheat, oat, and pea grown under either conventional or organic agriculture across farms in New York State (USA). Distinct microbiome clustering patterns indicated that organic and conventional production methods imposed strong selective pressures, shaping microbial assemblages within each group more distinctly than site or plant species variations. Using SCRS-based microbial phenotyping, we identified distinct microbial adaptations in agricultural soils, with organic systems favoring lipid-accumulating phenotypes for energy storage and stress resilience in low-input environments, while higher nutrient availability in conventional systems promoted carbon-rich phenotypes, enhancing rapid carbon assimilation and biomass production. Through network analysis of ecological hub species, we identified *Pseudomonas*, a plant growth-promoting rhizobacteria (PGPR), along with several nitrogen-fixing prokaryotes as core members within conventional agricultural systems. In contrast, organically managed soils featured PGPR taxa from the Bacilli class and contained microorganisms carrying antibiotic resistance genes, potentially indicating the presence of antibiotic resistance genes within organic agricultural environments. Overall, we found that the novel inclusion of microbial phenotyping methods, such as SCRS, can describe unique linkages between microbiome structure and their physiology that are distinctive between conventional and organic agricultural systems.

**Importance:** Our study successfully integrated single-cell Raman microspectroscopy and amplicon sequencing, two established techniques for analyzing microbial communities and their functions, enabling a link between genotype and phenotype to better characterize ecosystem dynamics. While few studies have explored microbial phenotypes alongside community composition to infer agricultural management differences, our research offered key insights into functional relevance of microbial communities to agricultural practices, demonstrating how management strategies influenced microbial adaptation. These findings advance microbial ecology research, demonstrating how agricultural management strategies influence microbiome structure and function, reinforcing the importance of phenotypic profiling in sustainable agriculture.

## 3 Introduction

The Green Revolution of the 1960s helped boost agricultural productivity with the widespread use of synthetic fertilizers, pesticides, and fuels to maximize crop yields globally. Decades after, conventional agricultural practices became widespread, but many have raised significant environmental concerns, including soil erosion, water pollution, pesticide overreliance, and biodiversity loss (1–7). In contrast to conventionally managed systems, organic agriculture typically limits synthetic fertilizer and pesticide use while promoting soil health practices for long-term sustainability (1, 3, 4, 6). Although organic systems are known to produce lower yields compared to conventional counterparts, they have ecological benefits that include less fertilizer input, enhanced floral and faunal biodiversity, and improved soil health in terms of soil aggregate stability, soil microbial biomass, the activities of dehydrogenase, protease, and phosphatase, and nutrient cycling (1). Thus, organic agriculture is considered a form of low-input food production that enables the conservation of soil, water, energy, and biodiversity to benefit long-term sustainable food production (3).

Understanding how different agricultural systems shape soil microbiomes is essential for evaluating their long-term effects on ecosystem resilience, nutrient cycling, and sustainable food production. Soil microorganisms play a vital role in agricultural productivity by forming complex networks of mutualistic, competitive, deleterious, and neutral interactions within interconnected ecological communities (8, 9). A community of microorganisms (microbiome) in a given agricultural soil habitat can shape soil ecosystem functions as a whole, extensively mediating soil nutrient cycling and nutrient supply for plants, soil moisture control, organic matter formation, soil remediation, and plant pest and disease suppression (4, 5, 10). Comparing soil microbiomes between conventional and organic systems reveals how farming practices influence microbial diversity, soil health, and nutrient cycling, informing sustainable agricultural strategies that balance productivity with ecosystem resilience.

Conventional and organic agricultural practices can influence soil microbial community composition and function due to their distinctive impacts on the soil environment. Many attempts have been made to compare soil microbiome characteristics between conventional and organic agriculture. Previous studies, spanning both short-term and long-term investigations, have documented significant differences in microbial community structure and functional traits between conventional and organic agricultural systems (4, 6, 11–15), but some sites showed only a minor difference in terms of microbial composition (11). In general, the distinctions between conventional and organic agriculture can arise from the differential use of agrochemicals such as chemical fertilizers and pesticides, among other practices. A review of research suggested that the overuse of pesticides had negative effects on soil microbial communities that in the worst case, led to significant yield loss, decreased soil microbial biomass, and reduced soil respiration (16). In contrast, organic agricultural soils are known for having greater microbial diversity and taxonomic heterogeneity compared to conventionally managed fields with crop rotations of wheat, barley, potato, lily, carrot, and maize (6) while Wang et al. (14) observed an increase in bacterial richness under organic systems. These findings suggest the potential benefits of organic farming in maintaining soil microbial diversity, which can contribute to improved ecosystem stability, nutrient cycling, and long-term agricultural sustainability.

However, the majority of comparative studies on soil microbiomes are constrained to one or more plant species or specific site variations, making it difficult to identify broad, overarching patterns across conventional and organic farming systems. For instance, many comparative studies on soil microbiomes in conventional and organic agriculture limited the number of sites and plant species analyzed to reduce confounding spatial, temporal, and plant variability (4, 6, 11–14, 17, 18), with the exception of a few large-scale surveys, such as the study of Chen et al. (15) that examined diverse vegetable plants grown in 30 conventional and organic greenhouses across China. While some research reported clear distinctions in microbial composition and biodiversity metrics (6, 14), both Chen et al. (15) and Jung et al. (13) detected no significant differences in alpha diversity of soil microbiomes between conventional and organic agricultural systems for greenhouse vegetables such as tomato, cucumber, eggplant, green pepper and rice. The lack of consistent differences in biodiversity across studies suggests that broader environmental variables may play a significant role in shaping soil microbial communities. Possibly, variability in soil biogeography, temperature, microclimatic factors, plants, and on-farm practices, likely contribute to challenges in interpreting impacts of conventional and organic agriculture on soil microbial diversity (4, 19, 20). To address these limitations, large-scale microbiome surveys incorporating diverse crops and geographic regions are essential for identifying robust and generalizable trends in microbial dynamics across conventional and organic farming systems.

Among the various approaches for comparing microbiome variations, single-cell Raman spectroscopy (SCRS) has recently emerged as a promising method for microbial characterization. While well-developed and widely applied genotyping techniques, such as 16S rRNA sequencing, have been used to compare genotypic variations, phenotyping the rhizomicrobiome has rarely been explored due to the lack of suitable techniques with sufficient resolution (21). Existing approaches in phenotyping the rhizomicrobiome either rely on specific biomarkers, such as metabolomics coupled with stable isotopes in tracing rhizomicrobial pathways (22), and/or only measure in bulk, such as enzymatic activity assays which only reflect the collective phenotypic profile of the rhizomicrobiome (23). The SCRS, as a label-free phenotyping method, offers a non-invasive, high-resolution metabolic fingerprinting of the microbiome that reflects the cellular composition and proteomics-like metabolics state of microorganisms in the soil ecosystem (20). Previously, few applications of SCRS in environmental microbial analysis were mostly restricted within simpler matrices such as highly engineered enhanced biological phosphorus removal (EBPR) systems (24, 25), and/or by the reliance on labeled biomarkers such as the use of stable isotope probing in nitrogen fixation activity (25). To date, only a few attempts in characterizing specific groups of the microbes from the rhizomicrobiome with SCRS have been documented (25, 26).

In this study, we performed a large-scale comparison of soil microbiomes derived from conventional and organic agricultural systems spanning 12 different plant species grown across various sites in New York State, USA. Specifically, we examined the soil microbiomes of 10 conventionally managed plants and 12 organically managed plants grown in different farm locations. We characterized different compositions of soil prokaryotic microbial communities between conventional and organic agriculture using the 16S rRNA sequencing. In complement to the commonly used amplicon sequencing, we employed a SCRS method to pioneer the characterization of phenotypic profiles of the rhizomicrobiome of various crops grown in conventional and organic farming systems, using a similar method to our previous study (26) to enable us to obtain single-cell-resolution phenotypic profiles of rhizomicrobiome in various crops that reveal functional information beyond taxonomic units. Combining this SCRS with amplicon sequencing, we aimed to link phenotypic characteristics with genetic variations, resulting in a deeper insight into microbial functions and interactions within conventional and organic farming systems. To summarize, our objectives were to: 1) identify distinct attributes of soil microbial communities in the rhizosphere of 12 different plant crops grown in conventionally and organically managed agricultural systems, and 2) link the complex microbial structure-phenotype relationships within the rhizomicrobiome under the influence of different agricultural practices.

## 4 Results

### 4.1 Distinct rhizomicrobial taxonomic structures across crop plants between conventional and organic agriculture

We obtained 1,709,785 ASVs with a mean of 15,544 ASVs per sample. A total of 4,151 bacterial species were discovered and belonged to 33 phyla, 267 families, and 460 genera. The phylogenetic relationships and compositional variations in the soil microbiome were examined across multiple taxonomic levels, comparing organic vs. conventional conditions across all plant species. (Supplementary Figure S1). Most species were distributed across seven bacterial phyla, Acidobacteriota, Actinobacteriota, Bacteroidota, Chloroflexota, Gemmatimonadota, Proteobacteria, and Verrucomicrobiota, with others belonging to other phyla that make up less than 0.5% of the total abundance. At the family and genus levels, soil microbiomes displayed considerable diversity, with a broad spectrum of taxa shaping overall composition across different plant species and production systems. These distinct shifts, driven by variations in soil type, plant species, and crop management practices, highlight the complexity captured in our large-scale comparative study, enabling the identification of general trends across conventional and organic systems.

Non-metric multidimensional scaling (NMDS) ordinations based on Bray-Curtis dissimilarity demonstrated clear differences in microbial community composition between conventional and organic systems, forming two distinct 95% confidence ellipses (permutational analysis of variance (PERMANOVA), permutation = 999, R^2^ = 0.92, the Benjamini-Hochberg (BH)-adjusted *P* < 0.05) (Figure 1A). Microbial communities within each agricultural system—conventional or organic— formed distinct clusters across all plant species, resulting in clearly separated groupings between the two systems. This suggests that farm production system (organic vs. conventional) plays a more dominant role in shaping microbial community structure compared to plant species and soil type, despite samples being collected from different locations and soil environments. In other words, the clear clustering patterns indicate that organic and conventional production methods exerted strong selective pressures, overriding environmental variability and driving distinct microbial assemblages. Furthermore, microbial communities were influenced by plant species, both within and across production systems, reflecting the selective role of host plants in shaping their associated microbiomes (PERMANOVA, permutations = 999, R² = 0.92, BH-adjusted *P* < 0.05) (Figure 1B). Notably, pairwise PERMANOVA analysis revealed distinct microbial profiles across all plant species between conventional and organic systems, highlighting the broad and consistent influence of farming practices on microbiome composition. Additionally, Supplementary Figure S2 illustrates that plant taxonomic variation—at both the family and species levels—played a significant role in shaping soil microbial community structure, as revealed by beta diversity patterns and supported by pairwise PERMANOVA analyses. These findings indicate that, beyond production style, plant taxonomy was also a key determinant of rhizosphere microbiome composition. PERMANOVA results, including R² values, raw *P* values, and BH-adjusted *P* values, are summarized in Supplementary Table S1. Overall, this result emphasizes the dominance of farming practices in structuring soil microbial communities, with plant species acting as an additional but secondary driver of microbiome composition.

**Figure 1.**
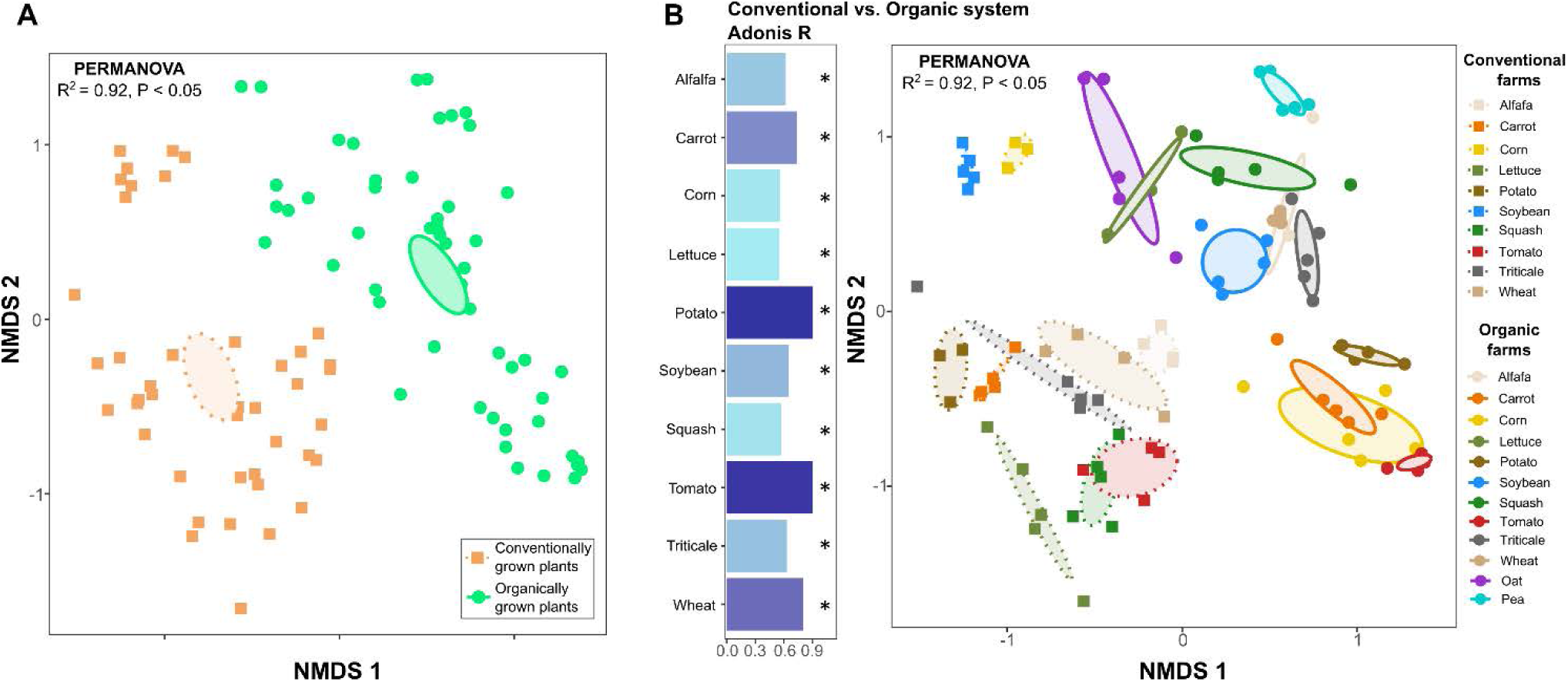
Non-metric multidimensional scaling (NMDS) ordinations based on Bray-Curtis dissimilarity illustrating variations in microbial community structures. (A) PERMANOVA analysis revealed a significant difference in microbial community structure between conventional and organic agricultural systems, with the statistical results displayed in the upper left corner of the plot. (B) Community composition varied based on plant species and production practices, as indicated by the PERMANOVA results presented in the upper left of the plot. Statistical significance between conventional and organic systems for each plant was assessed using pairwise PERMANOVA, with Adonis R² and BH-adjusted *P* values displayed on the left side of the NMDS plot (*: *P* < 0.05). A 95% confidence ellipse was generated for each plant group, with colors assigned to match their respective categories.

When analyzing microbial community structures across conventional and organic production systems using differential abundance analysis using the R package ALDEx2 (27), certain microbial phyla emerged as uniquely linked to each system despite differences in plant species and environmental conditions. Overall, Bacteroidota was more prevalent in the rhizosphere soils of conventional agriculture, whereas Proteobacteria, Firmicutes, Actinobacteriota, and Verrucomicrobiota were more abundant in the organically managed rhizomicrobiome (Figure 2A). This was further evidenced by pairwise comparison revealing the consistently higher relative abundances of Bacteroidota in the rhizosphere of conventionally grown carrot, soybean, squash, and triticale when compared to those in organic systems. Conversely, organic agriculture favored Proteobacteria (alfalfa, with minimal CLR difference), Firmicutes (carrot, corn, lettuce, potato, squash, tomato, and triticale), Actinobacteriota (alfalfa, lettuce, squash, tomato, triticale, and wheat), and Verrucomicrobiota (alfalfa, lettuce, potato, soybean, triticale, and wheat) (Figure 2B). These findings highlight the distinct microbial phyla signatures associated with conventional and organic agricultural systems, demonstrating that farming practices can apply meaningful selective pressures on soil microbial communities. ALDEx2 results, showing differentially abundant microbial phyla across plant samples, are presented in Supplementary Table S2, including CLR-transformed abundance differences, effect sizes, and BH-adjusted *P* values.

**Figure 2.**
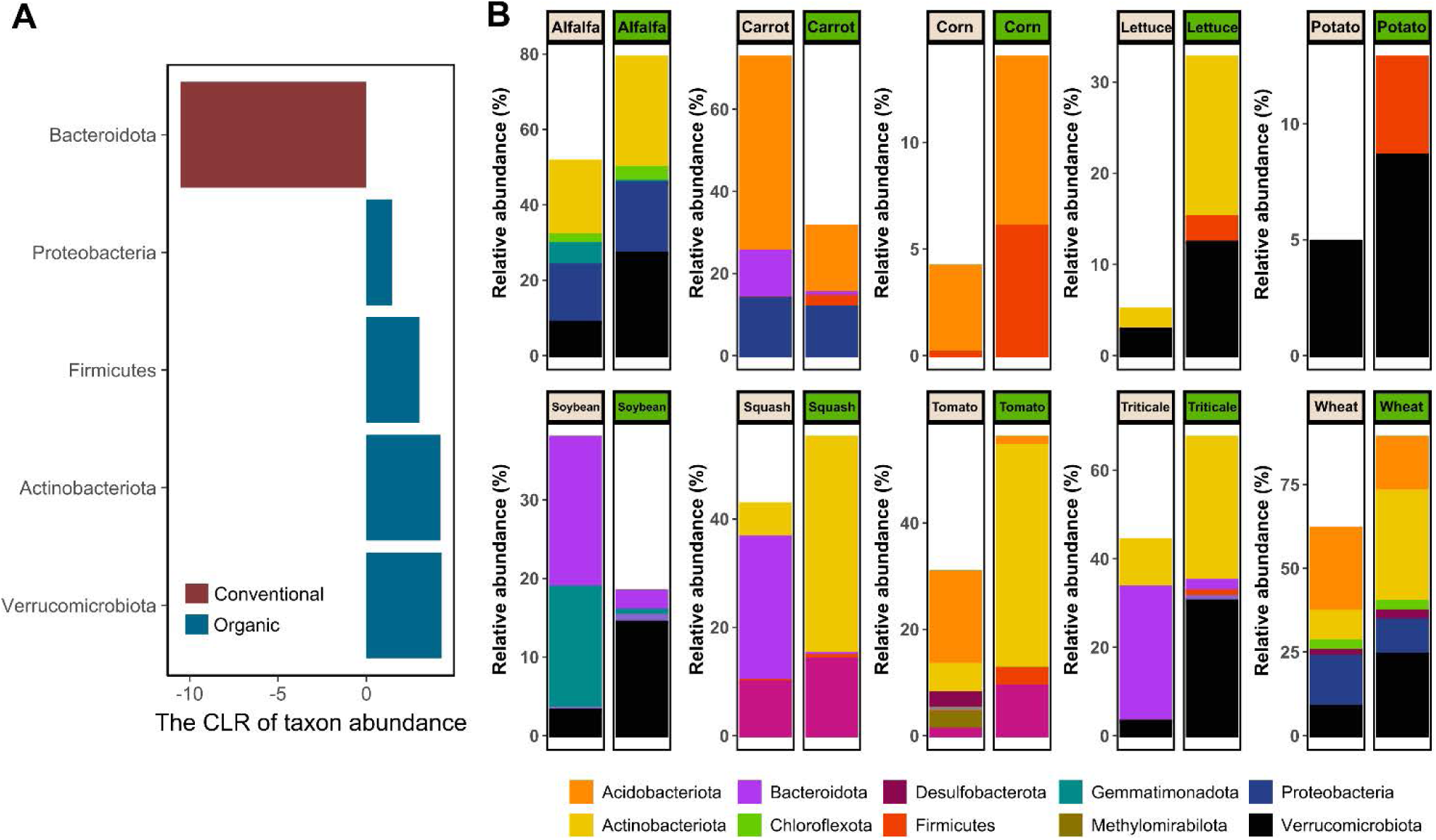
Differential abundance of microbial phyla in the soils between conventional and organic production using ALDEx2. (A) General patterns in microbial phyla with distinct relative abundances between conventional and organic agriculture, presented using centered log ratio (CLR) transformed abundances. Only phyla with significant differences in overall comparisons between the two systems are included, with red bars indicating higher abundance in conventional systems and blue bars representing higher abundance in organic systems. B) Bar graphs exhibiting variations in microbial phyla distribution across plant comparisons within the two agricultural systems. Statistical significance was assessed using pairwise Wilcoxon rank-sum test, at the BH-adjusted *P <* 0.1. Only phyla showing significant differences between conventional and organic systems are displayed, color-coded according to their respective microbial classifications.

The average distance to the centroid was used to compare the heterogeneity of the soil microbial community (Figure 3A). The distance to the centroid measures how closely each sample aligns with the cluster’s center, where smaller distances indicate tight clustering and high similarity, while larger distances reflect greater variability and higher heterogeneity within the group. Overall, the heterogeneity of beta diversity was not significantly different between conventional and organic agriculture (permutation = 9999; *F* = 0.76, permuted *P* = 0.39), indicating that both production systems showed similar levels of heterogeneity in soil microbial communities. Using alpha diversity indices such as Shannon diversity and species evenness, we observed higher microbial diversity in conventional alfalfa, squash, tomato, and wheat compared to their organic counterparts, whereas lettuce showed greater Shannon diversity under organic management (Figure 3B). For several other plant species, no notable differences were observed between conventional and organic systems.

**Figure 3.**
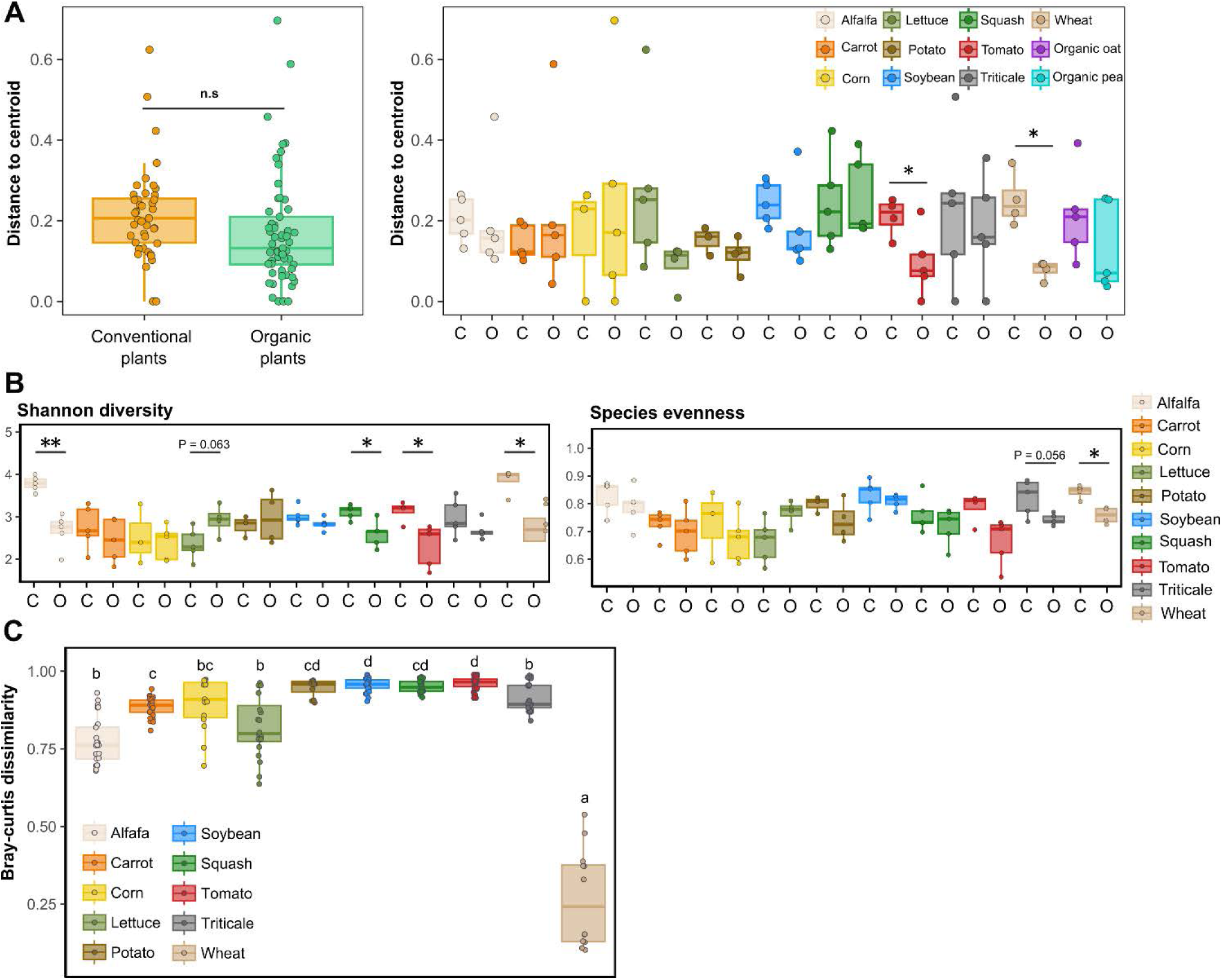
Heterogeneity and beta diversity of soil microbial communities. (A) The comparison of beta dispersion of soil microbial communities between conventional and organic agriculture. Pairwise comparison of the beta diversity dispersion for the same plant species between the production systems was performed at permutation number = 9999, *P* < 0.05. Statistical results were evaluated using a t-test, which yielded no significant differences, indicated by the label n.s. (not significant). (B) Shannon diversity and species evenness of each sample were compared between the two agricultural systems. (C) Pairwise comparison for Bray-Curtis dissimilarity of soil microbial communities between conventional and organic agriculture. Pairwise comparison of diversity indices and Bray-Curtis dissimilarity was performed using pairwise Wilcox rank sum test at the Benjamini-Hochberg adjusted *P <* 0.1. Significant *P* values were shown as asterisks (specific *P* values were written for *P* ≤ 0.1; *: *P <* 0.05; ** : *P <* 0.01). Box plots were composed of the center line indicating the median; the bottom and upper limits, the 25th and the 75th percentiles; and the whiskers extended 1.5 times of the interquartile range from the 25th and 75th percentiles. Abbreviations were as follows: n.s, no significance; C, conventionally grown plants; O, organically grown plants.

These findings suggest that microbial diversity varied across plant species, but neither conventional nor organic systems consistently showed higher alpha diversity or structural heterogeneity in soil microbial communities. The absence of significant differences in overall microbial community diversity indicates that farming practices did not substantially impact microbial variability at a broad scale.

Despite similar overall microbial community diversity between conventional and organic agriculture, most plant species had high Bray-Curtis dissimilarity scores, indicating distinct microbial community structures between the two systems when analyzed individually for each plant (Figure 3C). Soybean and tomato exhibited the greatest differences in compositional heterogeneity, whereas wheat showed only a minimal median dissimilarity of 0.24 between conventional and organic systems. These findings suggest that while broad microbial diversity and heterogeneity trends were comparable across agricultural systems, individual plant species experienced notable compositional shifts influenced by farming practices. Supplementary Table S3 provides detailed Bray-Curtis dissimilarity values for both pairwise comparisons of plants between conventional and organic systems as well as comparisons within each production group.

### 4.2 Key microbiome hub species revealed from network analysis of conventional and organic agricultural soils

Additionally, we identified hub species within the microbial networks of conventional and organic systems, based on their placement in the top 10% for normalized betweenness centrality and degree of connectivity (*P* < 0.1) (Supplementary Figure S3), following the approach described by Agler et al. (28). Betweenness centrality represents a taxon’s function as a connector, facilitating interactions among microbial members, while degree of connectivity is related to its ecological importance based on extensive associations within the community (29, 30). Together, these metrics help pinpoint key microorganisms that play a structural and functional role in shaping microbial communities, referred to as hub species. These hub species function as important ecological components, contributing to the stability and sustainability of microbial community structure (28, 31). Their high connectivity with other taxa and central positioning within the network help define ecosystem dynamics and stability. In ecological terms, the loss of a hub species can lead to the alteration or disruption of the whole microbiome system (29, 30).

Using SparCC (32), we constructed microbial networks representing microbiomes from conventional and organic agriculture, referred to as the conventional network and organic network throughout the text. We identified 483 soil microorganisms involved in 7,505 significant interactions within the co-abundance network of microbiomes associated with conventionally grown plants (Figure 4A). Within the conventional network, three hub microbial species were identified, including members of the genus *Rudaea*, genus *Nitrososphaera*, and class RBG-16-71-46. To further examine their role, we constructed a co-abundance network specifically linked to these hub species (hereafter referred to as hub network), which encompassed 769 co-abundance interactions with 66 microbial taxa (Figure 4B). The limited number of hub species in the conventional network indicates that hub taxa formation in conventional systems was potentially highly selective, shaped primarily by environmental pressures rather than broad microbial interactions. Factors such as agrochemical applications and tillage likely imposed constraints, influencing which microbial taxa could establish central roles within the network (32). Supplementary Table S4 provides details on co-abundance network interactions within the conventional hub network, including linked members, SparCC correlation coefficients (R), and *P* values. The strongest interactions in the hub network of rhizomicrobiomes in conventional systems were identified between members of the genus *Palsa-739* and the family Gemmatimonadaceae (R = 0.66), followed by the positive correlation between the class RBG-16-71-46 and the order Bacillales (R = 0.57); by the genus *Rudaea* and the genus *CF-154* (R= 0.56); by the family Steroidobacteraceae and the family Polyangiaceae (R = 0.55); by the genus *Mesorhizobium* and the genus *VAYN01* (R = 0.53); and by the genus *Rudaea* and the genus *Ginsengibacter* (R=0.52).

**Figure 4.**
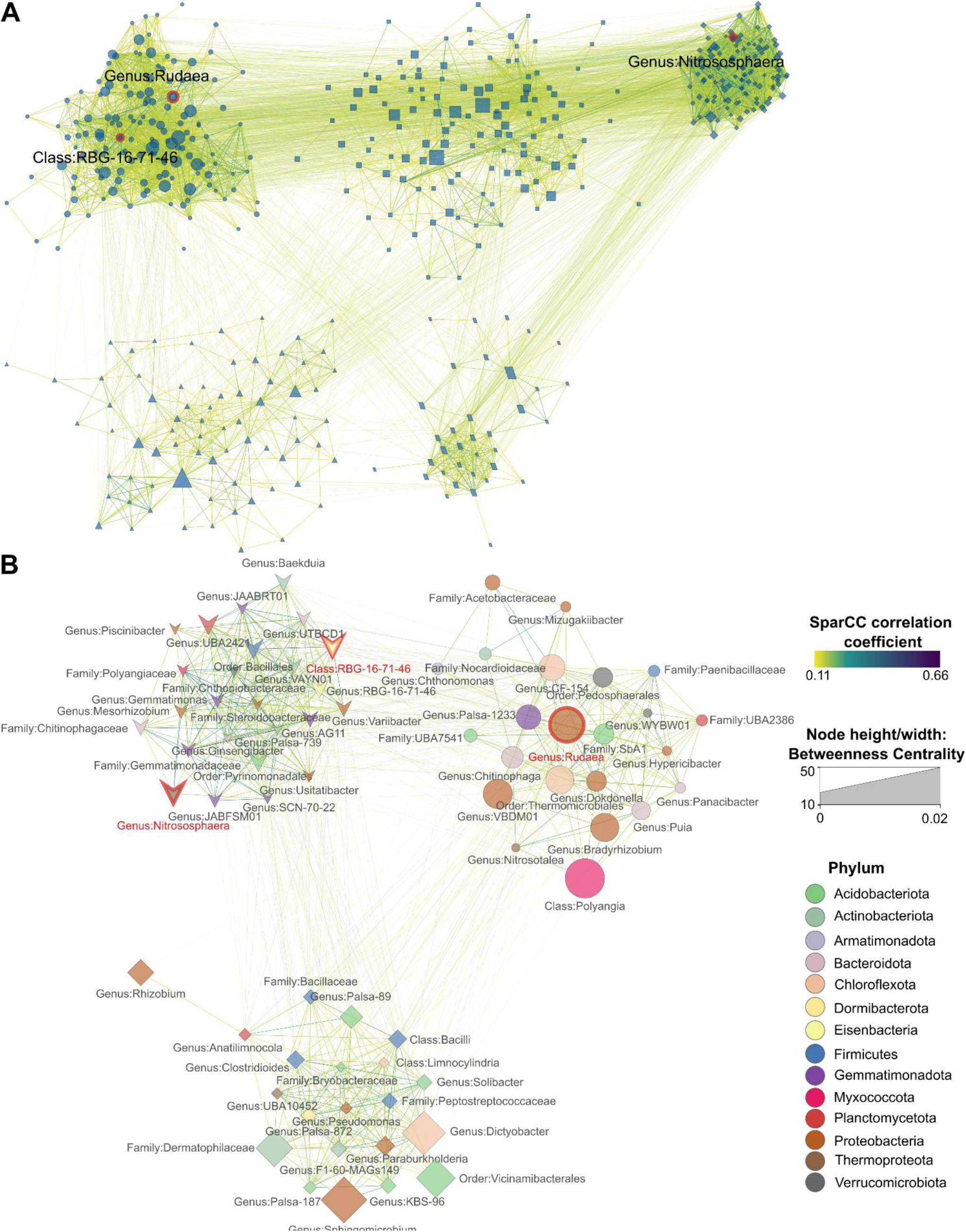
Microbiome SparCC co-abundance networks. (A) Co-abundance network of the whole rhizomicrobiomes from conventional agriculture. (B) The co-abundance network of the hub species in conventional farming systems. Hub species are annotated at their lowest taxonomic level and highlighted with red border lines and red text. Nodes represent individual taxa identified at their lowest taxonomic level, with node colors corresponding to their respective phyla. Node size reflects the betweenness centrality of each taxon, indicating its importance in microbial interactions. Edges connecting nodes are color-coded to represent their correlation strength. Only statistically significant correlations (pseudo *P* < 0.1, derived from bootstrapping with 100 repetitions) were included in each network. Clusters were generated using the Markov Clustering Algorithm (MCL), with each cluster’s nodes depicted in different shapes to distinguish groupings within the network.

We observed 348 microorganisms involved in 3,168 co-abundance interactions in the rhizosphere of plants grown under organic agriculture (hereafter referred to as the organic network) (Figure 5A). In the organic network, we discovered eight hub microbial taxa such as the genus *JABFSM01*, the genus *Tumebacillus*, the family Hyphomicrobiaceae, the genus *Caulifigura*, the genus *UTCFX2*, the genus *Bacillus*, the family Chitinophagaceae, and the genus *Paenibacillus*. In the hub network of microbiomes in organic agricultural systems, 41 microbial taxa were found to engage in 217 significant interactions with the eight hub species (Figure 5B). The presence of eight hub species in the organic network—more than twice the number found in conventional systems— suggests that organic farming, devoid of agrochemicals and extensive tillage, may reduce selective pressure, allowing a broader range of hub taxa to be shared among organically grown plants rather than favoring a few dominant species. These shared ecological characteristics among organically grown plants may contribute to the stability of hub networks, ensuring resilience even amid environmental fluctuation (33). A detailed summary of all co-abundance interactions in the organic hub network, including linkage members, SparCC correlation coefficients (R), and *P* values, is provided in Supplementary Table S5. In the organic hub network, the strongest interactions were found between species associated with the genus *Microvirga* and the genus *Paenibacillus* (R = 0.42), followed by the genus *VBCG01* and the family Burkholderiaceae (R = 0.37); by the genus *Paenibacillus* and the genus *Ensifer* (R=0.35); by the genus *Mycobacterium* and the genus *Solirubrobacter* (R = 0.31); by the genus *Paenibacillus* and the family Burkholderiaceae (R = 0.31); and the genus *Thermoactinomyces* and the genus *Paenibacillus* (R = 0.31).

**Figure 5.**
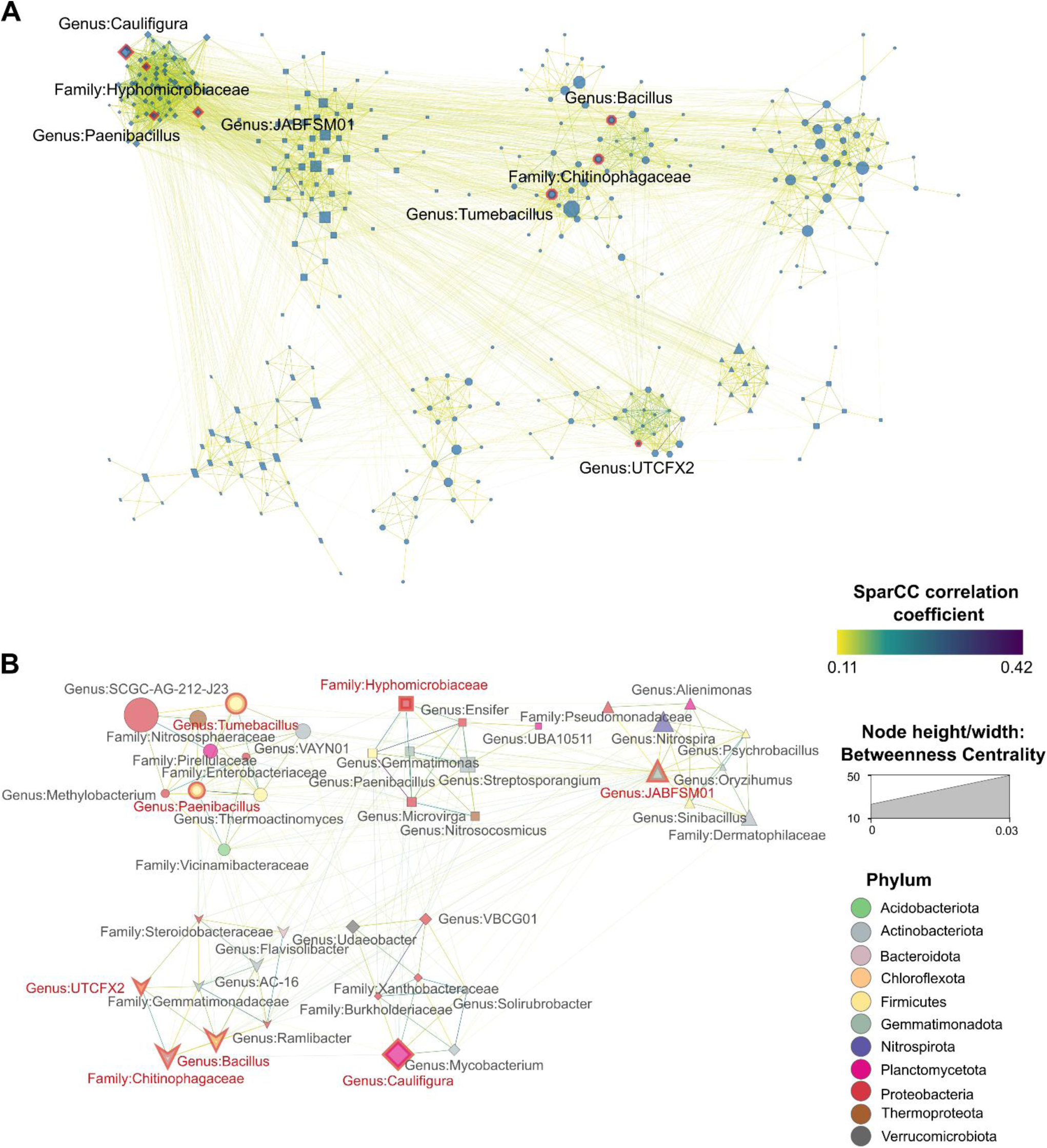
Microbiome SparCC co-abundance networks. (A) Co-abundance network of the whole rhizomicrobiomes from organic agriculture. (B) The co-abundance network of the hub species in organic farming systems. Hub species are annotated at their lowest taxonomic level and highlighted with red border lines and red text. Nodes represent individual taxa identified at their lowest taxonomic level, with node colors corresponding to their respective phyla. Node size reflects the betweenness centrality of each taxon, indicating its importance in microbial interactions. Edges connecting nodes are color-coded to represent their correlation strength. Only statistically significant correlations (pseudo *P* < 0.1, derived from bootstrapping with 100 repetitions) were included in each network. Clusters were generated using the Markov Clustering Algorithm (MCL), with each cluster’s nodes depicted in different shapes to distinguish groupings within the network.

### 4.3 Single-cell Raman fingerprints and phenotype clustering of microbiomes associated with different plants

We used SCRS to conduct single-cell phenotypic profiling of the rhizomicrobiome across various crops, generating 3,285 Raman spectra, each representing the metabolic fingerprint of an individual cell. While operational taxonomic units (OTUs) are defined by nucleotide sequence similarities, clustering microorganisms based on genetic markers (e.g., 16S rRNA gene for bacteria) to infer taxonomic relationships, OPUs are derived from phenotypic similarities observed in single-cell Raman spectra captured by SCRS. These spectral profiles reflect biochemical composition and metabolic activity, allowing for functional grouping independent of genetic sequence data. Hierarchical clustering grouped 66% of the collected spectra into 21 OPUs, while the remaining 34% were singletons with little similarity to any other spectra. The distinctive distribution of the identified OPUs of rhizomicrobime samples of all plants are shown in Figure 6A. Among the identified OPUs, OPU-0 and OPU-1 were both ubiquitous, occurring in all samples regardless of crop type or farm operation style, and abundant, collectively accounting for half of all acquired spectra (Figure 6A). The other OPUs were less abundant and not universally present among all plants. No significant differences (*P* > 0.05) were found in terms of the overall OPUs abundance distribution or the abundances of specific OPUs between conventional and organic plants.

**Figure 6.**
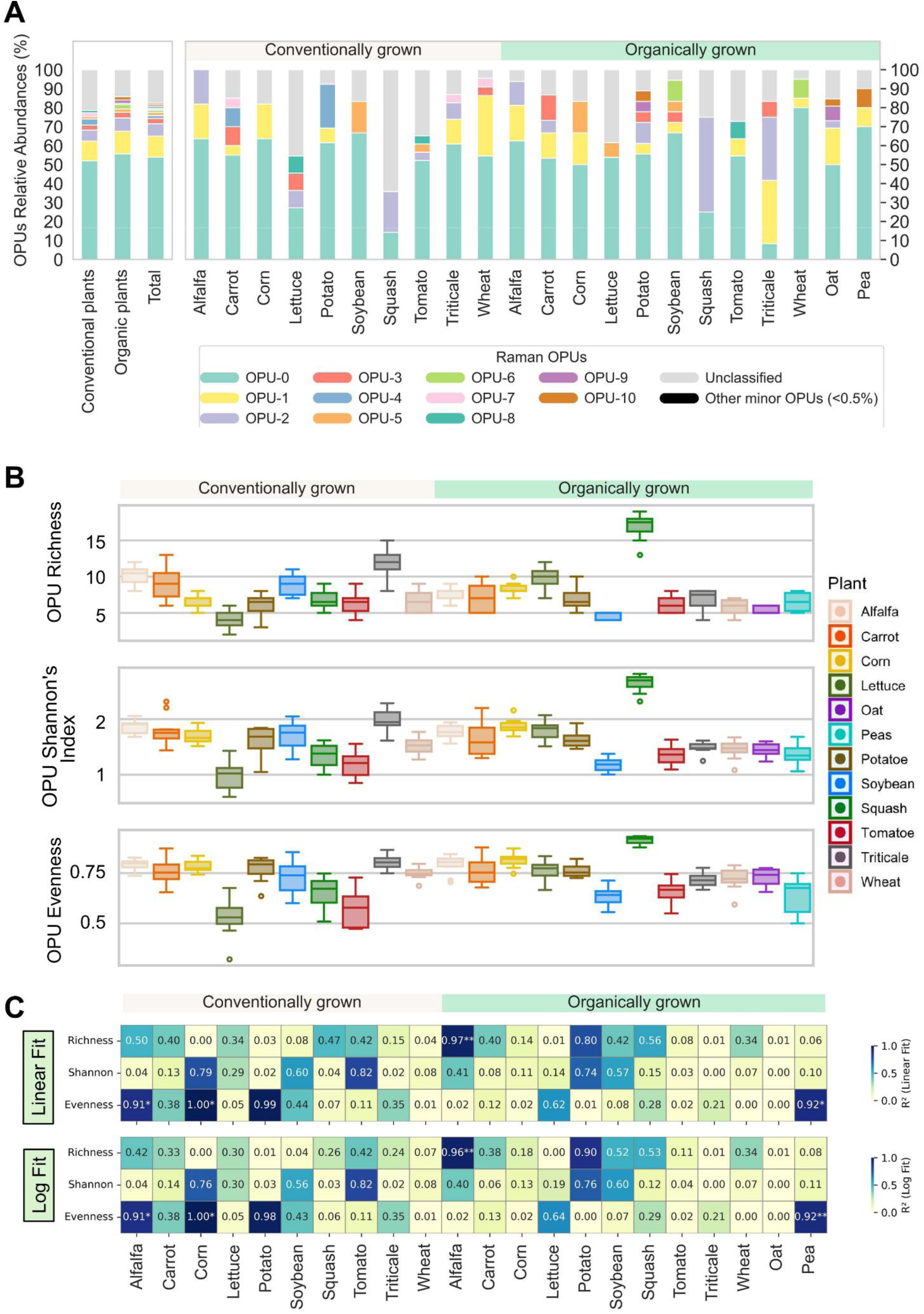
Characterization and comparison of Raman profiles of rhizomicrobiomes of conventionally and organically managed plants. (A) Phenotypic profiling of the microbial community based on operational phenotypic units (OPUs) distribution among different crops surveyed. The left panel shows the OPUs distribution for conventional plants, for organic plants, and overall, while the right panel displays those for each individual crop. A total of 21 OPUs are identified and numerically labeled sequentially and only 12 OPUs with relative abundances over 0.5% are shown, while the other 9 minor OPUs are combined as “Other minor OPUs (< 0.5%)”. Raman spectra that did not get classified into the 21 OPUs are combined as “Unclassified”. (B) Distribution of richness, Shannon’s diversity index, and evenness of OPUs. (C) Heatmaps showing the coefficient of determination (R^2^) from linear and log-transformed linear regression models between ASV-based (taxonomic) and OPU-based (phenotypic) alpha diversity metrics across samples, revealing phenotype-genotype correlations. Each cell represents the R^2^ value for a given diversity metric (richness, Shannon, or evenness) within a treatment, fitted between ASV and OPU values. Significance is indicated by asterisks: *: *P <* 0.05; ** : *P <* 0.01; and ***: *P* < 0.001.

Distinct Raman fingerprints could also be identified for each OPU, underscoring substantial variations in their cellular molecular compositions. Leveraging the Fisher rank score analysis and the correspondence between Raman shifts and certain biomolecular bonds or structures, especially in nucleic acids, protein, carbohydrates, and lipids (34), the metabolic and physiological implications of observed discrepancy in Raman spectra can be further elucidated (Supplementary Figure S4). For example, OPU-0 and OPU-1, though both prevalent, exhibited distinctly different Raman features. OPU-0 showed characteristic peaks around 1250 and 1450 cm^−1^, linked to CH_2_ structures in lipids (34), whereas OPU-1 was characterized by peaks at 1358 and 1574 cm^−1^, associated with amino acids and peptide bonds in proteins. OPU-8 phenotype was highly concentrated with the Raman spectra in ranges between 600-800 cm^−1^ related to peptides and proteins (tyrosine, amide IV, amid V, and tryptophan), nucleotides (guanine, adenine, uracil, thymine, cytosine), and cytochrome C (34). Among these, OPU-8 was most enriched at 687 cm⁻¹ (23.1%), associated with trimethylsilyl (Si-CH) rocking vibrations (35), suggesting enhanced carbon uptake and metabolism in this phenotype (35).

The beta diversity of the microbial phenotypes associated with different plants was further elucidated by the SCRS-metabolites profiles as OPUs (Figure 6B). Consistent with previous observation in the heterogeneity from ASVs (Figure 3B), the richness, Shannon’s index, and evenness were not significantly different between conventional and organic plants (*P* > 0.05). However, for specific plants, variations in the phenotypic structures between the soil microbial communities in conventional and organic farms showed that conventional lettuce and squash had lower richness, Shannon’s index, and evenness than their organic counterparts, although many other plants did not exhibit a notable difference between conventional and organic systems. To further assess the relationship between taxonomic and phenotypic diversity, we performed linear and log-log regression analyses between ASV- and OPU-derived diversity metrics (Figure 6C). Strong and significant couplings between taxonomic and phenotypic diversity indices were observed in conventional alfalfa and corn as well as organic alfalfa and pea, and the linear-transformed models seemed to yield higher or at least equivalent R^2^ values. These results potentially indicated a low functional redundancy as the phenotypic diversity had not plateaued yet scaling to taxonomic diversity.

Soil microbiomes were strongly associated with soil functionality, as evidenced by the distinct distribution of abundant OPU communities across conventional and organic systems. (Figure 7A). Figure 7A shows that 35.5% of the variation in microbial community structure was explained by distinct OPU members, emphasizing their influence in shaping the functional divergence of rhizosphere microbiomes under conventional and organic agricultural systems. We applied Distance-based Redundancy Analysis (dbRDA) to assess how microbial community structures correlated with OPU distributions, uncovering key microbial phenotypic traits that distinguished conventional and organic agricultural systems, with statistical validation using PERMANOVA (R² = 0.19, F = 4.54, *P* < 0.001). This approach underscores the distinct microbial assemblage shifts driven by differing agricultural practices. Several OPUs such as OPU-1, OPU-3, OPU-8, OPU-10, and unclassified OPUs were significantly correlated with the variations of microbiomes (Envfit, permutation = 10,000, *P* < 0.1). The direction of each OPU arrow in the dbRDA ordination space indicates the strength and direction of correlation between the OPU and the site points within the ordination configuration (36). Among the significant OPUs, conventional microbiomes were strongly associated with OPU-8, OPU-10, and unclassified OPUs, whereas organic microbiomes showed strong correlations with OPU-0 and OPU-3.

**Figure 7.**
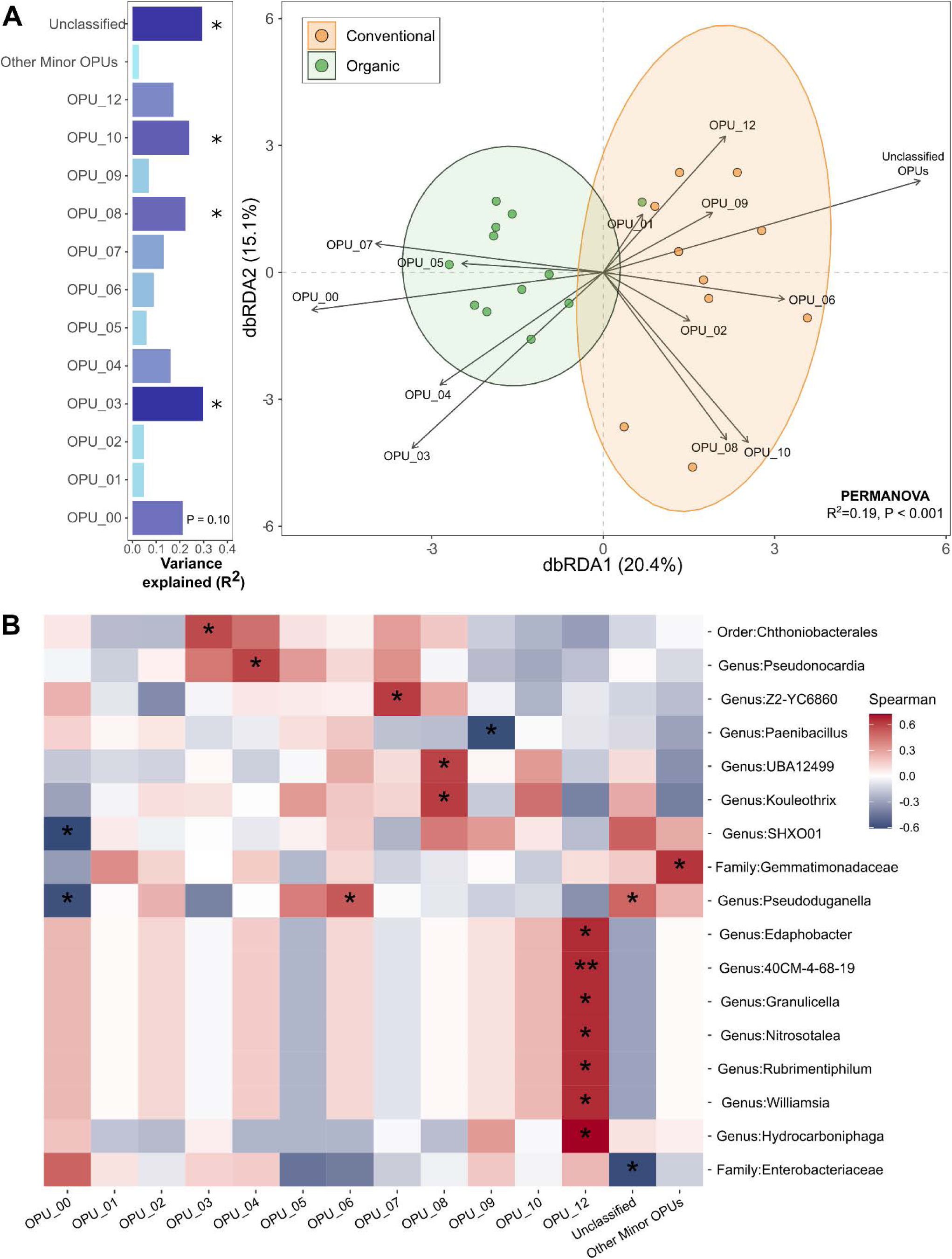
Microbiome variability and functional interactions in conventional and organic production systems. (A) Distance-based Redundancy Analysis (dbRDA) examining the relationship between microbial community composition and OPU distributions, with microbial phenotypes treated as environmental variables driving microbiome differentiation between conventional and organic soils. The variance explained by each OPU was assessed using the EnvFit function in the vegan R package and depicted in horizontal bar graphs positioned to the left of the dbRDA plot. Asterisks indicate significant OPUs, showing functional groups that were differentially associated with microbial communities in conventional and organic systems at *P* < 0.1. (B). Heatmap showing Spearson correlations between microbiome structure and OPU abundance. Significant microbial taxa were identified at the BH-adjusted *P* < 0.1.

To further explore microbial-functional associations, we conducted Spearman correlation analysis, identifying significant microbial taxa linked to OPU abundance at the BH-adjusted *P* < 0.05 (Figure 7B). Among OPUs significantly associated with rhizomicrobiomes in organic systems, OPU-0 exhibited negative correlations with species belonging to the genera *SHXO01* and *Pseudoduganella*, while OPU-3 showed a positive association with a species of the order Chthoniobacterales. For conventional microbiomes, OPU-8 displayed significant positive correlations with members of the genera *Kouleothrix* and *UBA12499*, highlighting distinct microbial-functional associations between the two agricultural systems. Several OPUs which formed notable correlations with specific microbial genera included OPU-4 which was correlated positively with genus *Pseudonocardia* and OPU-6 which exhibited a positive relationship with genus *Pseudoduganella*. Similarly, OPU-7 was positively linked to genus *Z2-YC6860*, whereas OPU-9 showed a negative correlation with genus *Paenibacillus*.

## 5 Discussion

### 5.1 Similarities and dissimilarities in soil microbial community structure between conventional and organic agriculture

The most notable finding of our study is that microbial community structures in conventionally managed farms showed similarities across plant crop types, likewise, microbiomes in organic farms showed similarities across organically cultivated plants. The distinct clustering patterns emerged within their respective agricultural systems, despite variations in plant species and site conditions. This observation offers a broader ecological perspective, extending insights beyond single-plant analyses within a single site. Unlike many studies that focus on a single plant or a few plants at the same site, our research took a regional-scale approach, comparing 10 sets of plant species (alfalfa, carrot, corn, lettuce, potato, soybean, squash, tomato, triticale, and wheat) in conventional versus organic systems, plus two additional crops (oat and pea) in organic systems. Despite soil and plant variability, we observed intriguing patterns, including an increase in Actinobacteriota, Firmicutes, and Verrucomicrobiota and a decrease in Bacteroidota, Gemmatimonadota, and Acidobacteria in organically cultivated plants. These findings align with previous studies, although trends cannot be generalized among all plant species. For instance, other researchers reported the high abundances of Actinobacteriota and Firmicutes in the soils of organic cereal crops (12), finger millet managed through regenerative agriculture (37), and organically grown tomato that were suppressive against tomato corky root disease (38); Firmicutes in soils under long-term management as organic systems (4, 11, 17); and Verrucomicrobiota in organic tea tree cultivation (39). These findings highlight the significant impact of agricultural management on soil microbial communities, highlighting distinct structural patterns between organic and conventional systems.

Other microbial taxa showed notable community profiling in the different farming systems. Copiotrophic bacteria, known for their ability to flourish in nutrient-rich conditions, showed clear distinctions between organic and conventional agricultural systems (40). Copiotrophic microorganisms increased in both organic and conventional systems, with Firmicutes and Actinobacteriota more abundant in organic farms, while Bacteroidota showed higher levels in conventional systems, reflecting distinct microbial adaptations to agricultural conditions. Previous studies have also observed similar trends, showing that the increased abundance of Firmicutes in organic systems was associated with the use of composted manure and organic matter, which created a nutrient-rich environment that supports their growth (41, 42). Similarly, many species of the Actinobacteriota are known to prosper in environments with high soil organic matter and nutrient availability (38, 43, 44). A meta-analysis of 56 peer-reviewed studies, encompassing 149 pairwise comparisons across diverse climatic zones and experimental durations (ranging from 3 to over 100 years), reported that organic systems had 32% to 84% higher microbial biomass carbon, microbial biomass nitrogen, total phospholipid fatty acids, and enzymatic activities (dehydrogenase, urease, and protease) compared to conventional systems (45). These findings indicate that organic farming enhances nutrient availability, which may contribute to the proliferation of copiotrophic microorganisms such as Firmicutes and Actinobacteriota. Conversely, Bacteroidota were more abundant copiotrophic bacteria in conventional agricultural systems, as reported in previous studies (11, 46). Their abundance in conventionally managed soils is likely due to the nutrient-rich conditions created by synthetic fertilizers, which support their proliferation. In this context, both conventional and organic agriculture provided nutrient-rich environments, as reflected in the abundance of copiotrophic bacteria. However, variations in microbial composition may result from distinct nutrient preferences, with Firmicutes and Actinobacteria thriving on organic matter-derived nutrients, while Bacteroidota may preferentially utilize nutrients directly from fertilizers. Collectively, both conventional and organic systems supported copiotrophic bacterial growth, but variations in nutrient sources and farming inputs likely shaped microbial community composition and adaptation patterns differently.

Our study revealed a high abundance of Verrucomicrobiota in the rhizosphere of organic agricultural plants compared to conventional ones. These enigmatic microorganisms are well-adapted to nutrient-poor environments, efficiently utilizing available resources in ways that enable their survival where many other microbes struggle (8, 47, 48). Particularly, their ability to process complex organic compounds makes them to play a crucial role in soil carbon cycling, contributing to nutrient turnover and long-term soil health (49–51). The low-input nature of organic agriculture, with its gradual nutrient release, may create ideal conditions for these bacteria. Studies suggest that organic soils treated with low-rate organic fertilizers (0.7 LU per hectare) tended to maintain lower carbon and nitrogen levels, favoring the growth of oligotrophic microorganisms such as Verrucomicrobiota (52). The gradual nutrient release in organic farming—regulated by composted manure, plant residues, and organic fertilizers—may generate localized nutrient-rich zones that promote copiotrophic bacterial growth while simultaneously sustaining low-nutrient pockets that favor oligotrophs. This dynamic nutrient distribution enables diverse microorganisms to thrive under resource-limited conditions, aligning with our findings. Consequently, organic agriculture supports the growth of these versatile microbial groups, which play significant roles in nutrient cycling and soil health.

### 5.2 Co-abundance patterns reflecting ecosystem characteristics and farming practices

Co-abundance networks analyze microbial community patterns by detecting taxa that reliably co-occur across samples, uncovering ecological relationships such as mutualistic interactions or shared environmental dependencies (4). In the organic network, we found several hub species belonging to Bacilli class members such as the genera *Tumebacillus*, *Bacillus*, *Paenibacillus*, and *Brevibacillus*, which are known as plant growth promoting rhizobacteria (PGPR) for their beneficial traits such as nutrient mineralization and disease control (53–55). Our findings align with multiple prior studies that have reported high abundances of these Bacilli species in organic agricultural soils (4, 12, 56, 57), further supporting the strong association between organic amendments and Bacilli community dynamics. Additionally, we discovered several microorganisms capable of carrying antibiotic-resistant genes (ARGs), including species belonging to the genus *Mycobacterium*, the family Enterobacteriaceae, the family Vicinamibacteraceae, the family Burkholderiaceae, and the family Steroidobacteraceae (57–59). The presence of ARG-associated microorganisms within the hub network of microbial communities in organic agricultural soils may be linked to the frequent application of animal manures, a practice known to contribute to the spread of ARGs (60). This practice may accelerate the spread of ARGs in organic agricultural systems, contributing to the emergence of antibiotic-resistant bacteria. The presence and distribution of these microbial hosts in organic agricultural soils raise ongoing concerns regarding their potential impact on microbial ecosystems and resistance dissemination.

In conventional agricultural systems, our study identified numerous beneficial microorganisms within their hub network, despite the widespread use of agrochemicals such as pesticides and synthetic fertilizers. These include key microbial groups involved in nitrogen fixation (e.g., *Rhizobia*, *Bradyrhizobia*, *Mesorhizobium*, and *Paraburkholderia*) (61–63), nitrogen cycling (e.g., *Nitrososphaera*) (64, 65), and plant growth promotion (e.g., *Pseudomonas*) (66). Surprisingly, nitrogen fixers were more active for microbial networking in fields managed with synthetic fertilizers than the organic hub network. This increased activity may stem from higher phosphorus availability, aligning with findings from Orr et al. (67), who reported elevated *nifH* gene activity—a key gene involved in nitrogen fixation—in fields managed with synthetic fertilizers compared to those receiving compost amendments. They concluded that phosphorus enrichment through synthetic fertilization enhanced nitrogen-fixing microbial performance, offering insights into the complex relationship between nutrient inputs and microbial dynamics in agricultural systems. Additionally, xenobiotic-degrading microorganisms such as *Pseudomonas* and *Sphingomicrobium* played an active role in the conventional hub network. These organisms are known for breaking down polycyclic aromatic hydrocarbons (PAHs), potentially thriving on PAHs introduced into soils through chemical pesticides and herbicides (68, 69). The presence of xenobiotic-degrading microorganisms in the conventional hub network suggests that the frequent input of agrochemicals possibly led to the enrichment of xenobiotic microbial members that can degrade and mineralize these chemicals (70). Despite the extensive application of agrochemicals in conventional systems, beneficial microorganisms—such as nitrogen fixers, nutrient cyclers, PGPR, and xenobiotic degraders— continued to play a crucial role in microbial networks.

These findings suggest that both conventional and organic agricultural systems were able to form beneficial microbial communities in their hub network, though the composition and functions of these microbial members were greatly influenced by distinct management practices. Linking microbial networks to agricultural strategies enhances our understanding of ecosystem dynamics, offering valuable insights for developing practices that support soil health and long-term sustainability.

### 5.3 Linking microbial phenotypes to agricultural practices using a SCRS-based approach to microbiome analysis

When microbial diversity indices did not show significant differences between organic and conventional agriculture, SCRS can still provide valuable insights by characterizing the phenotypic and metabolic profiles of microbial communities (25, 71). By analyzing the biochemical composition of individual microbial cells, SCRS enables the identification of functional dynamics and interactions that diversity indices alone may overlook. In comparing organic and conventional agricultural practices, distinct microbial phenotypes were associated with each system. Organic agricultural soils were enriched with the phenotypes of lipid-accumulating microorganisms (OPU-0) which are possibly efficient in lipid accumulation owing to enhancing conversion of substrate to triacylglycerols or polyhydroxyalkanoates (PHAs), fatty acid synthesis, and intracellular lipid storage (72). This finding is consistent with our observations of increased growth of Actinomycetota and the actinomycete genus *Mycobacterium* (a key member of the organic hub network) in organically managed soils. Actinomycete species possess the capability for *de novo* fatty acid biosynthesis from acetyl-CoA and can store substantial amounts of triacylglycerols, sterol esters, or long-chain lipid mycolic acids (C_60-90_) within their cells (72–75). The ability to store intracellular lipids benefits the survival of these microorganisms during starvation periods, as lipids serve as an important energy substrate, aiding in cell growth, division, and stress resilience under nutrient-limiting conditions (72). Thus, the lipid-accumulation phenotype likely improves the resilience of Actinomycetota in organic systems, where nutrient availability fluctuated due to the gradual release of nutrients from organic matter.

Conversely, conventional soils showed higher associations with a carbon-rich phenotype (OPU-8), such as those involving the genus *Kouleothrix* and a species belonging to the family UBA12499 and phylum Methylomirabilota. These microorganisms are known to be efficient in carbon metabolism and storage, with *Kouleothrix* playing a significant role in carbon polymer hydrolysis and fermentation (76), and Methylomirabilota members (e.g. UBA12499) utilizing volatile fatty acids for methanogenesis (77). These metabolic processes supported microbial activity in conventional systems, enabling them to efficiently convert available carbon into energy, thereby sustaining immediate growth and functionality. Such phenotypes differed from those in organic systems, which favored lipid-accumulating microorganisms adapted for long-term energy conservation, whereas conventional systems were characterized by active metabolism and energy conversion. The frequent inputs of synthetic fertilizers and agrochemicals in conventional agricultural practices may ensure a steady supply of readily accessible nutrients, driving microbial activity toward faster energy extraction and storage.

Our study surveyed soil microbiomes in the rhizosphere of 22 crop plants under conventional and organic agricultural systems, revealing clear differences in microbial composition, network structures, and phenotypes, despite soil and plant variability. Additionally, our integration of SCRS with microbiome analysis offers a novel approach for linking microbial phenotypes with genotypes, advancing the ability to characterize functional diversity across conventional and organic agricultural systems. By revealing distinct microbial adaptations—such as lipid accumulation in organic soils and carbon-cycling phenotypes in conventional soils—our study lays the groundwork for future investigations into biochemical and molecular mechanisms underlying plant-microbe interactions relevant to sustainable agriculture.

## 6 Discussion

### 6.1 Farm surveys and sample collection

We sampled rhizosphere soils from a total of 12 commonly found and ecologically important crop plants, including potato, tomato, carrot, lettuce, squash, corn, wheat, soybean, alfalfa, triticale, oat, and pea from two conventional farms and two organic farms (Table 1). Figure 8 illustrates the spatial distribution of collection sites both within and between farms, highlighting variations in location, soil type, and environmental heterogeneity. The two organic farms, which adhered to restrictions on synthetic fertilizers and pesticides, while utilizing organic amendments such as animal manures, were certified in compliance with the regulations of the United States Department of Agriculture (78). In contrast, the two conventional farms applied agrochemicals, including chemical fertilizers, pesticides, and herbicides, while complying with federal and state regulations governing their use (79). Sampling was performed during the summer of 2021 (June 28^th^ to July 22^nd^) with a local temperature averaged at 20°C and an average local precipitation of 176.53 mm. For each plant, 5 replicated plants were removed at the time of sampling using a soil auger, and rhizosphere soils at the depth of 0-20 cm were carefully collected, and kept under 4 °C during the transportation to the lab. Soil samples were either processed immediately for single-cell Raman microscopy analysis, or stored under −80 °C until further analysis. The soil auger was meticulously cleaned and sterilized with 70% ethanol between sampling replicates to prevent cross-contamination.

**Figure 8.**
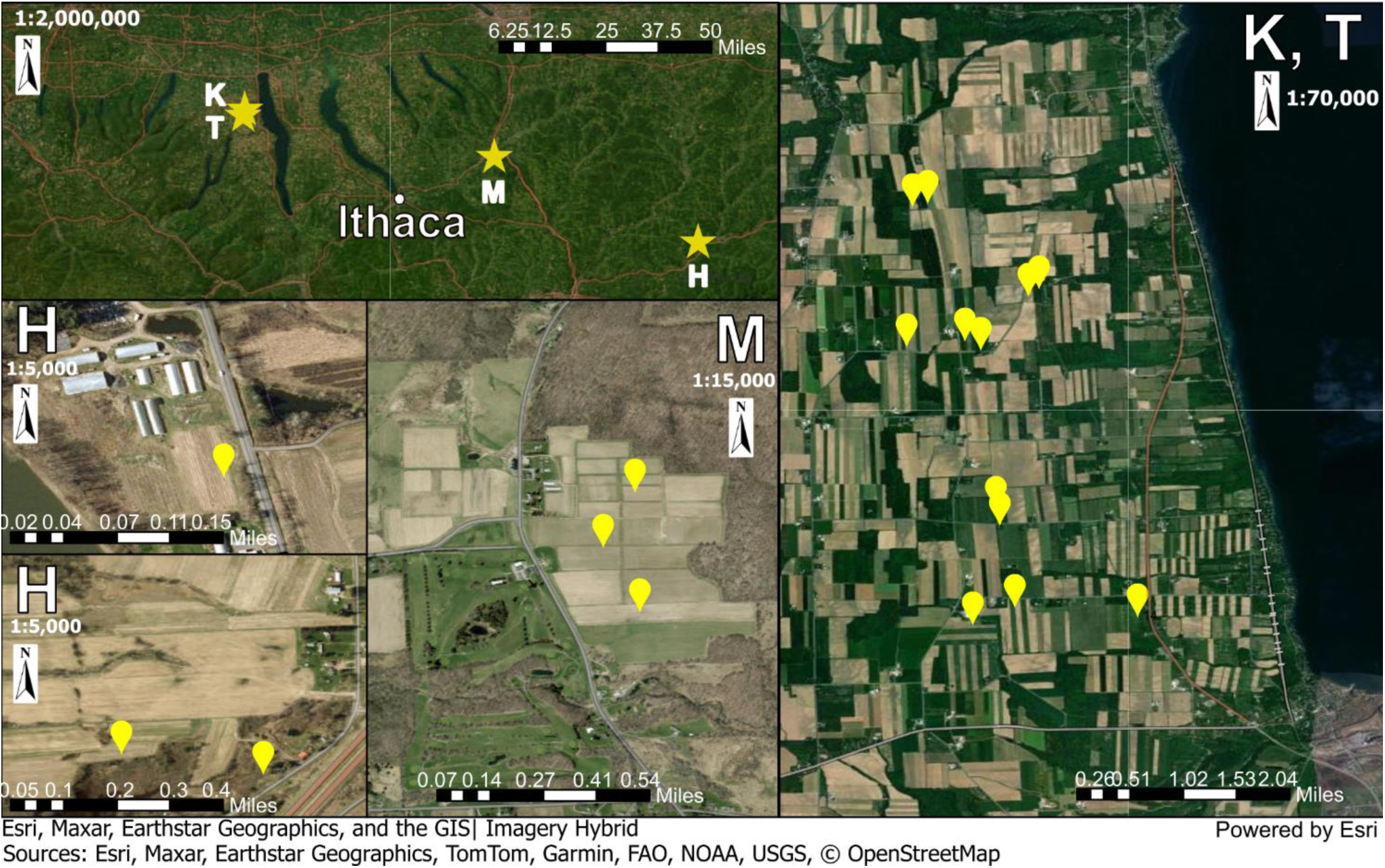
Farm locations of organic and conventional systems. Abbreviations of sites: M for Mainstreet farms, K for Lakeview Organic Grain LLC owned and operated by Klaas and Mary-Howell Martens, T for the conventional farm operated by Titus Zimmerman, and H for Hellers Farm CSA. The map is created by ArcGIS Pro 3.4 (Esri Inc., US) using the Imagery Hybrid map from ArcGIS Online (80).

**Table 1.** Summary of surveyed crop plants and source farms.

| **Crop name** | **Family** | **Scientific name** | **Conventional source farm** | **Organic source farm** |
| --- | --- | --- | --- | --- |
| Carrot | *Apiaceae* | *Daucus carota* | Hellers Farm CSA<br>(42°16'50.8"N<br>75°28'37.7"W) | Mainstreet farms<br>(42°34'5.0"N<br>76°11'34.2"W) |
| Lettuce | *Asteraceae* | *Lactuca sativa* |  |  |
| Potato | *Solanaceae* | *Solanum tuberosum* |  |  |
| Squash | *Cucurbitaceae* | *Cucurbita pepo* |  |  |
| Tomato | *Solanaceae* | *Solanum lycopersicum* |  |  |
| Alfalfa | *Fabaceae* | *Medicago sativa* | Titus<br>Zimmerman's<br>farm<br>(42°42'17.3"N<br>76°59'36.5"W) | Lakeview Organic<br>Grain LLC<br>(42°41'34.5"N<br>76°59'48.0"W) |
| Corn | *Poaceae* | *Zea mays* |  |  |
| Soybean | *Fabaceae* | *Glycine max* |  |  |
| Triticale | *Poaceae* | × *Triticosecale* |  |  |
| Wheat | *Poaceae* | *Triticum aestivum* |  |  |
| Oat | *Poaceae* | *Avena sativa* |  |  |
| Pea | *Fabaceae* | *Pisum sativum* |  |  |

### 6.2 DNA extraction, PCR amplification, and library preparation

Rhizosphere soil samples were sieved through mm mesh and ground thoroughly before DNA extraction. We used 0.25 g of a soil sample to extract soil DNA using DNeasy PowerSoil Pro Kits (Cat#: 47014; Qiagen, Carlsbad, CA) following the company’s manual. The extracted DNA was amplified by primers 515F (5’-GTGYCAGCMGCCGCGGTAA -3’) and 806R (5’-GGACTACNVGGGTWTCTAAT -3’) to target the V4 hypervariable region of bacterial and archaeal 16s rRNA. PCR reactions were conducted using 10 μl of 2 x AccuStart II PCR ToughMix (QuantaBio, Beverly, MA, United States), 1 μl of 10 μM forward and reverse primers with final concentrations of 0.5 μM, 6 μl of DNase-free PCR Water (MoBio, Carlsbad, CA, United States), and 2 μl of sample DNA. The PCR thermocycling conditions were as follows: 94°C for 2 min; 25 cycles at 94°C for 20 s, at 55°C for 20 s, and at 72°C for 30 s; with a final elongation at 72°C for 5 min. We used the method introduced in Howard et al. (81) to prepare sequencing libraries based on unique two-barcode index combinations (Nextera, Illumina), followed by normalization on clear 96-well plates. The pooled libraries were paired-end sequenced using the NovaSeq 6000 S4 (Illumina, San Diego, CA, USA) at Novogene (Durham, NC).

### 6.3 Bioinformatics

Raw forward and reverse reads were trimmed using Cutadapt to remove adaptor contaminants (82). Forward and reverse reads which had more than Phred offset score 20 were merged to create single consensus sequences using BBMerge (83) at the default setting. Bioinformatics analysis and classification of microbial communities from sequencing data were conducted following the methods described by He et al. (26). Amplicon sequence variants (ASVs) were created using DADA2 (84) in the denoise-single mode on the QIIME2 platform (85). Uchime algorithm was applied to correct for amplicon sequencing errors and identify and remove chimeric reads. ASVs were taxonomically assigned using a Naive-Bayes classifier trained on the Greengenes2 2022.10 database through the q2-feature-classifier plugin (86). The ASV data were transferred to R v.4.3.0 (87) and rarefied to the lowest read depth of any sample in the dataset using the function *rarefy_even_depth* in the phyloseq package (88). The completeness of rarefaction was assessed based on coverage-based rarefaction and extrapolation (R/E) curves with Hill number (q=1, Shannon diversity) using the package iNEXT.3D (89–91) (Supplementary Figure S5). These curves help estimate how well the observed diversity represents the true diversity by plotting species accumulation as sequencing depth (sample size) increases. The results demonstrated that all samples exceeded 99% coverage after rarefaction, preserving most observed microbial diversity and enabling reliable comparisons across agricultural systems with minimal ecological resolution loss.

### 6.4 Microbiome diversity assessments

For rhizosphere microbiome processing, we employed the phyloseq package (88) in R v.4.3.0 (87) to efficiently handle and analyze microbial community data. Initially, the rarified ASV counts were converted to percentage values, and taxa with a relative abundance below 0.5% in each plant group were excluded to reduce statistical noise. The dataset was further refined by excluding taxa present in fewer than 80% of replicated samples, ensuring a robust ecological representation. The Bray-Curtis dissimilarity matrix was constructed using the function *distance* in the vegan package (92) and visualized via NMDS with the function *ordinate* in the phyloseq package (88). The ASV abundance data were transformed into CLR using the microbiome package (93) before significant variations in beta diversity was assessed using PERMANOVA by employing the function *adonis2* in the vegan package (92). The CLR transformation was performed to mitigate mathematical distortions that can occur when proportional microbiome data is directly analyzed with Bray-Curtis or Hellinger methods, ensuring a more accurate representation of microbial community differences (94).

Alpha diversity indices, including Shannon diversity and species evenness, were computed from the filtered dataset (80% prevalence threshold) using the function *diversity* in the vegan package (92). The pairwise Wilcoxon rank-sum test was conducted to evaluate statistical significance among alpha diversity indices, with multiple testing corrections applied using the BH method at *P* < 0.05, implemented via the package rstatix (95). To assess community beta diversity dispersion, we applied the method introduced by Lupatini et al. (6), providing insights into differences in microbiome heterogeneity associated with organic and conventional agricultural systems. The ASV dataset, filtered by the 0.5% abundance threshold but not by the 80% prevalence filter, was used to maximize beta diversity dispersion and transformed into relative abundance for subsequent analyses. The dataset was subjected to permutated betadispersion analysis using pairwise Bray-Curtis dissimilarities, implemented via the function *betadisper* in the vegan package (92). Significance of the difference in the heterogeneity between conventional and organic agriculture was tested by a t-test using the function *permutest* in the package vegan (92). All plots were generated in R using ggplot2 (96) with significant differences marked with different alphabets.

### 6.5 Differential abundance (DA) analysis

We conducted DA analysis based on the non-rarified ASV data, as rarified microbiome data are often prone to increased Type-I and Type-II errors in DA analysis of microbiome data (88, 97). The ANOVA-like differential expression (ALDEx) analysis was chosen for the DA analysis due to its low false discovery rates compared to other DA methods (97–99). The ASV counts were CLR-transformed after Monte Carlo samples of Dirichlet distributions were calculated using the package ALDEx2 (v.1.30.0) (27) in R. Significantly differential taxa among treatments were identified by the BH-adjusted *P* values < 0.05 from the Wilcoxon rank sum test.

### 6.6 Network analysis

The SparCC method was employed to conduct a co-abundance network analysis of soil microbiomes discovered in the soils of the plants grown under conventional agriculture and organic agriculture. SparCC was deemed more appropriate for microbiome network analysis as it effectively reduces compositional bias and data sparsity—common challenges in microbiome datasets— allowing for more reliable correlation inferences regarding microbial abundance (100). SparCC was computed at default settings using the SparCC Python codes available at https://github.com/dlegor/SparCC. Non-rarified ASV counts were utilized to calculate correlation coefficients and pseudo *P* values < 0.1. To generate a network, only significant correlations with associated pseudo *P* values of *P <* 0.1 were included using the igraph package (101) in R. The network was transported and visualized in Cytoscape V.3.10.0 (102) using the R package, Rcy3 (103). Hub species in the networks of conventional and organic systems were identified as those falling within the top 10% of normalized betweenness centrality and degree of connectivity (*P* < 0.1), following the method outlined by Agler et al. (28).

### 6.7 Raman fingerprinting and phenotyping clustering of rhizomicrobiome associated with different plants

SCRS-based rhizomicrobiome phenotyping was performed for all the collected rhizosphere soil samples associated with different plants, with detailed protocol being demonstrated elsewhere (26). Briefly, soil samples were mixed with the detergent solution (3 mM sodium pyrophosphate, 0.5% Tween 20, 0.35% polyvinylpyrrolidone in PBS) and shaken for 30 min to detach the rhizo-microorganisms off the soil particles. The soil slurry was then placed on top of 80% Histodenz density-gradient solution (w/v) and subjected to centrifugation at 14,000 g under 4°C for 1.5 hours. The extracted and collected rhizo-microorganisms were then washed, diluted, and disaggregated by passing through 26-gauge needles repeatedly before being placed on top of the CaF_2_ Raman slide (Crystran Ltd., UK).

Raman instrumentation, setup, and data processing can be found in our previous publications (25, 104). A total of at least 100 Raman spectra with biological signature peaks at 1003 cm⁻¹ and 1657 cm⁻¹, corresponding to phenylalanine and amide I, respectively, were collected to represent the rhizomicrobial communities of each plant. OPUs, defined based on the similarity of Raman spectra to cluster and categorize microorganisms into different groups, were created through multivariate clustering as described previously (25, 26), except for the cutoff value adjusted to 0.44 according to Akaike information criterion (AIC) to account for the high diversity of the rhizomicrobiome. To evaluate the relationship between taxonomic and phenotypic alpha diversity metrics, we performed both linear and log-transformed linear regression analyses between ASV- and OPU-derived richness, Shannon, and evenness values. Ordinary least squares (OLS) regression models were fitted for each treatment using the statsmodels Python package (v0.14.1) (105). In the log-transformed models, both the independent and dependent variables were transformed using the natural logarithm of one plus the value (log1p) to accommodate zero values and improve numerical stability using Python Numpy (v2.2.4) (106). For each model, the coefficient of determination (R²) and corresponding *P* value were extracted to assess the strength and significance of the relationship. Heatmaps were generated using seaborn (v0.13.2) (107) to visualize R² values across treatments and diversity metrics, with statistical significance indicated by asterisks: *P* < 0.05 (*), *P* < 0.01 (**), p < 0.001 (***). The dbRDA biplot was generated to assess significant Spearman correlations between different operational phylogenetic units (OPUs) and microbial abundance, utilizing the microeco package (v.1.8.1) (108) in R. The direction and strength of each correlation of different OPUs with soil microbiomes were identified using the function *Envfit* (permutations = 999) in the package vegan (92). Notable correlations between the OPUs and individual microbial members were determined by *P* values < 0.1.

## 7 Data availability

The raw sequencing data for soil microbial community amplicon sequences is available in the National Center for Biotechnology Information Sequence Read Archive (NCBI SRA) under BioProject ID: PRJNA1263029. Additional data generated during this study can be obtained from the corresponding author upon reasonable request.

## 8 Acknowledgments

The authors extend their gratitude to Hellers Farm CSA, Mainstreet Farms, Titus Zimmerman’s farm, and Klaas and Mary-Howell Martens for their participation in this study. Additionally, gratitude is extended to Victoria L. Giarratano, Assistant Director of Agriculture, Food Systems, and Community Development at Cornell Cooperative Extension, for invaluable assistance in facilitating connections with both conventional and organic farmers. This study was supported by funding from award no. 2019-67013-29364 from the U.S. Department of Agriculture’s National Institute of Food and Agriculture, along with a Fulbright Program graduate research scholarship awarded to Y.S. (PS00298891).

---

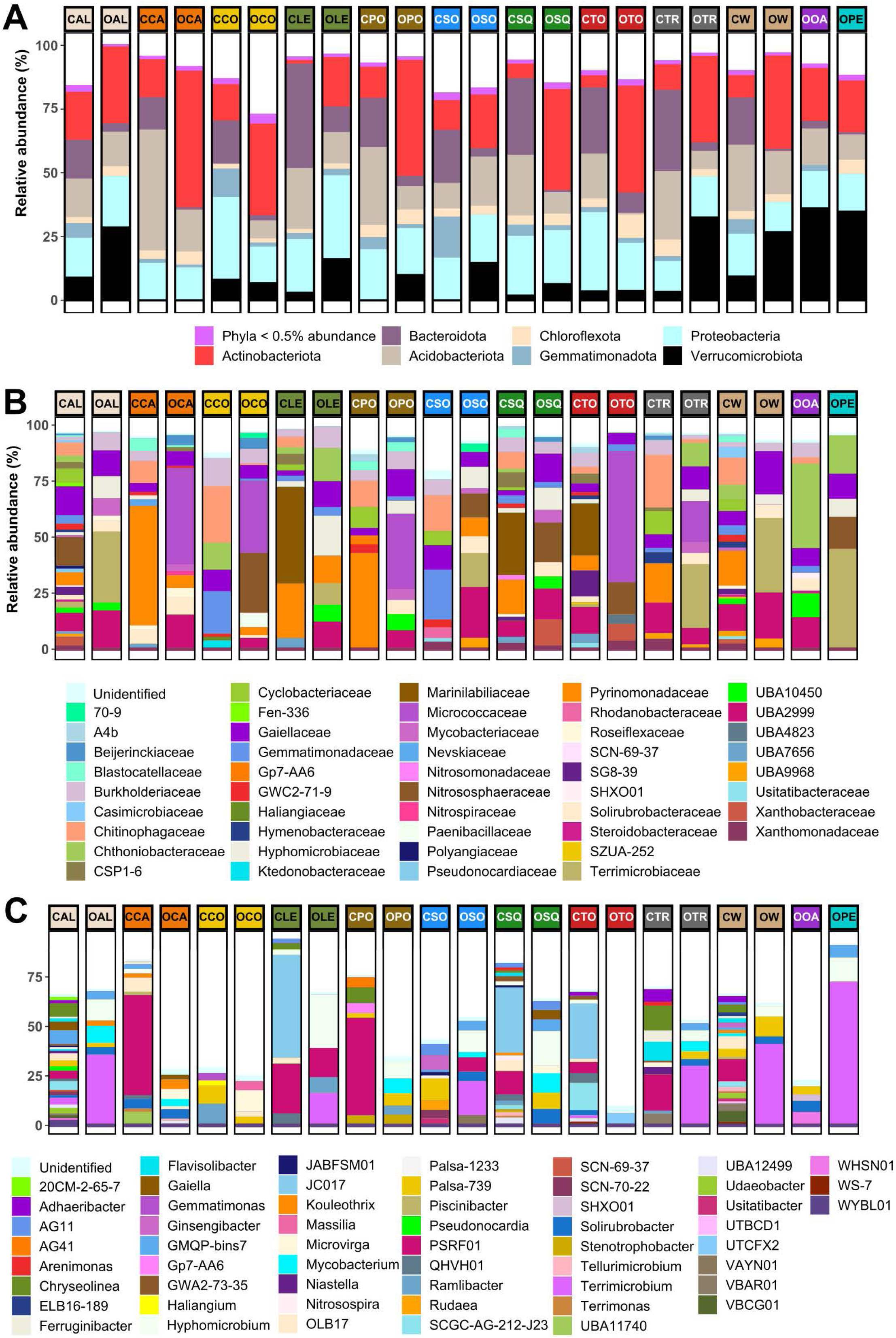

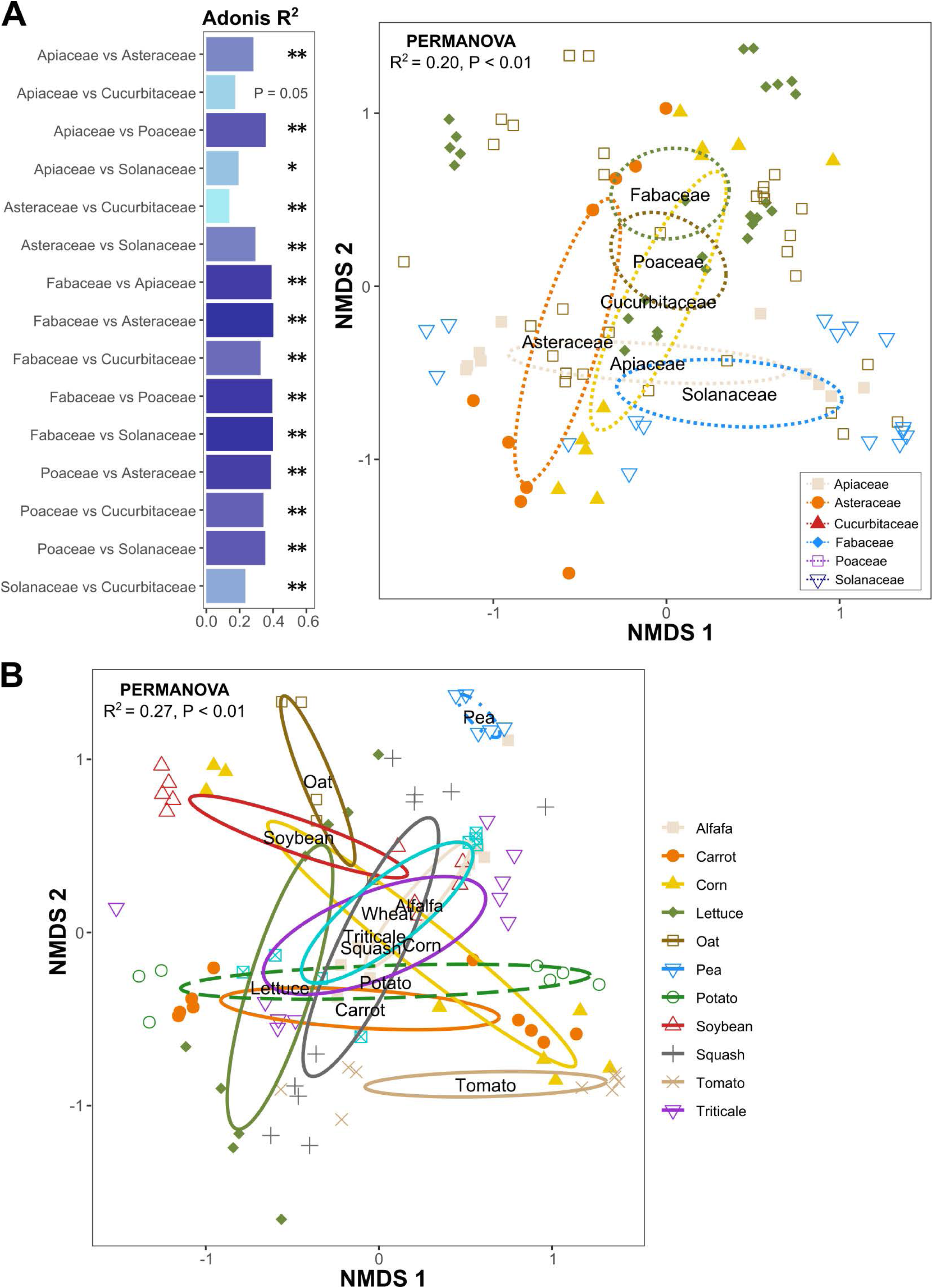

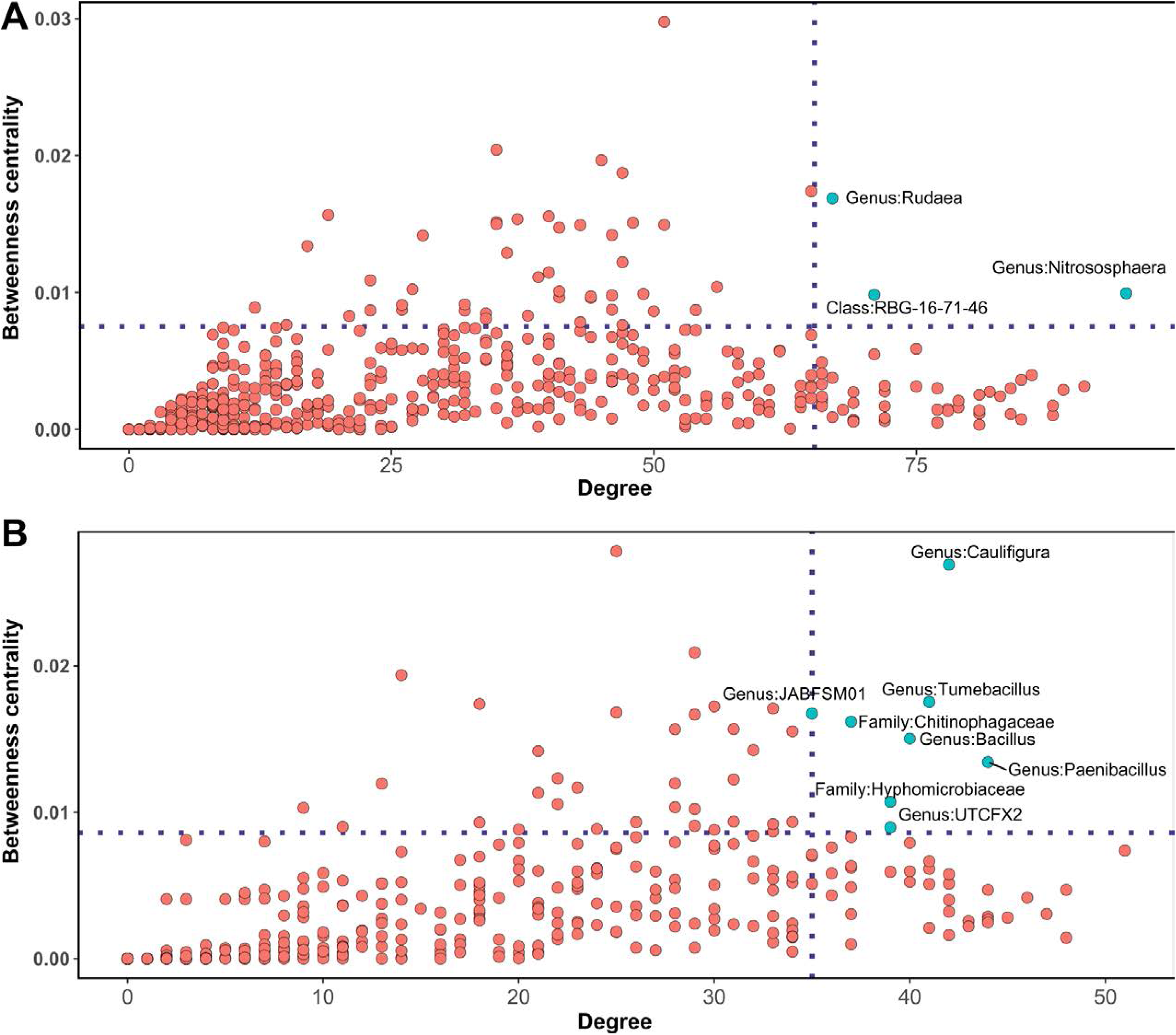

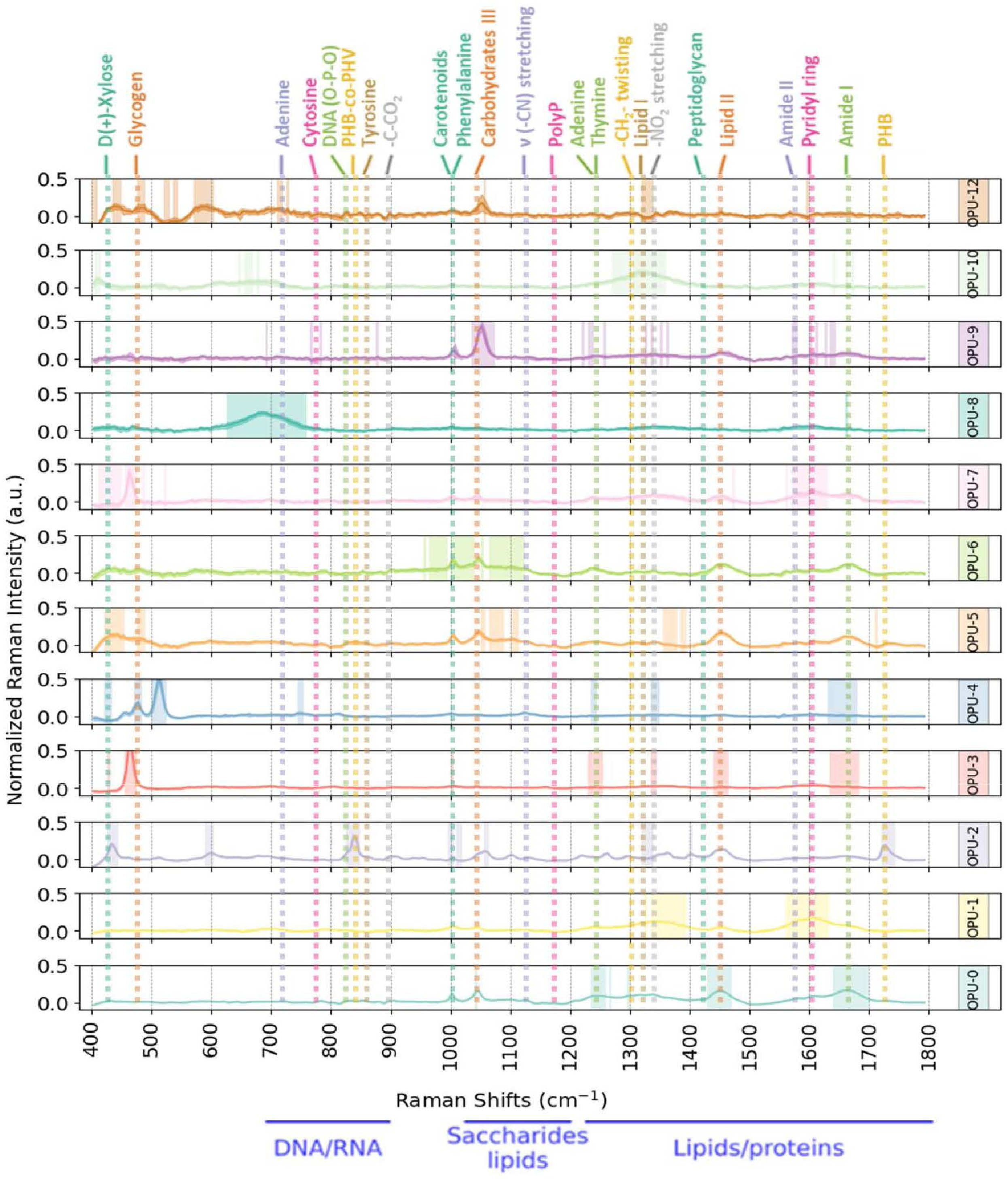

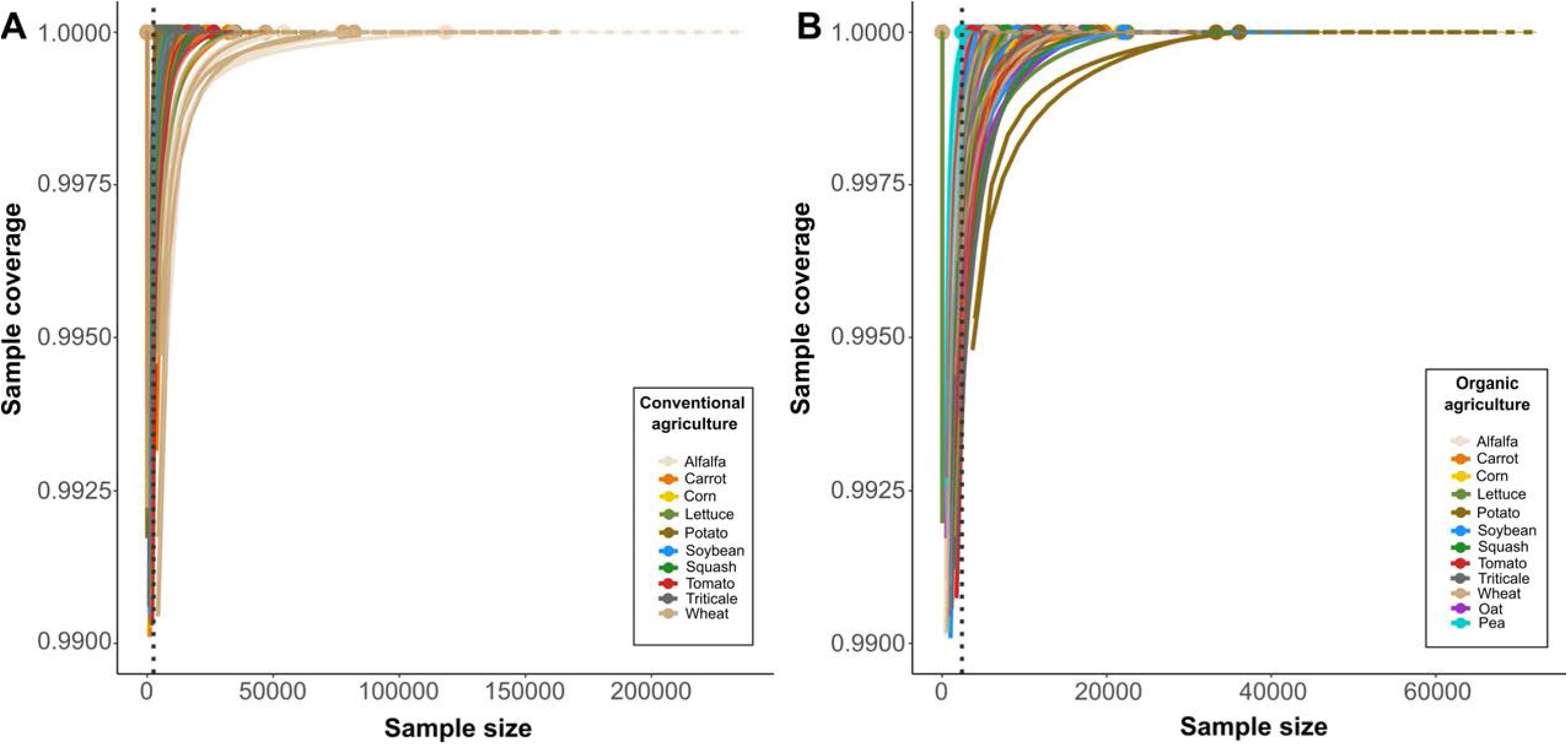

